# Microbes regulate glomerular filtration rate in health and chronic kidney disease in mice

**DOI:** 10.1101/2025.04.08.647647

**Authors:** Jiaojiao Xu, Eesha Verma, Jason Sanchez, Sepideh Gharaie, Sunyoung Jeong, Mahta Gooya, Kunal Gupta, Hamid Rabb, Jennifer L. Pluznick

## Abstract

Microbes are implicated in a variety of host physiological and pathophysiological processes. In this study, we tested the hypothesis that microbes modulate glomerular filtration rate (GFR). Microbiota were depleted in mice using oral antibiotics (ABX; a mixture of ampicillin, neomycin, and vancomycin). GFR was significantly increased in ABX-treated mice. To confirm that the increase in GFR was due to decreased microbes, we also measured GFR in germ-free (GF) mice. GFR was increased in GF mice as compared to both conventional and conventionalized GF (CGF) mice. We next used the murine adenine diet model to ask if suppressing gut microbes with ABX also increases GFR in a setting of chronic kidney disease (CKD), where GFR is impaired. In females on an adenine diet, ABX increased GFR versus adenine alone on weeks 4 and 6. In males, ABX elevated GFR on week 2. Adenine diet significantly increased plasma creatinine and kidney fibrosis; this was suppressed by ABX in both sexes. To explore the mechanism of this increase, we tested the hypothesis that altered tubuloglomerular feedback (TGF) contributes to elevated GFR using the sodium-glucose cotransporter 2 (SGLT2) inhibitor empagliflozin (EMPA); EMPA impairs Na^+^ reabsorption in the proximal tubule, altering TGF. We found that EMPA impaired ABX-induced GFR increases on week 3 but not week 5, suggesting that altered TGF contributes to the initial increase in GFR. In conclusion, the microbiome plays a key role in ‘setting’ baseline GFR by a mechanism which partially involves TGF, and, suppressing gut microbes can elevate GFR even in CKD.

**Translational Statement:** This study reports that GFR is elevated when gut microbes are absent or suppressed in mice, indicating a role for commensal microbes to help establish baseline GFR in health. Likewise, suppressing gut microbes also elevates GFR in a chronic kidney disease model. These data suggest a future possibility of modulating the commensal microbes to elevate GFR in a clinical setting.

## Introduction

Chronic kidney disease (CKD) is a major health issue with a global prevalence of 13%^1,2^. CKD is characterized by the progressive loss of kidney function^3^ which leads to disorders of electrolyte and acid-base balance, as well as anemia and bone disease. CKD is also strongly associated with an increased risk of cardiovascular events^2,4^ and hypertension^5^. Therapy to manage kidney disease includes blood pressure control and blockade of the renin angiotensin system^1^, as well as sodium-glucose cotransporter 2 inhibitors^6,7^. Despite these efforts, in a subset of patients, glomerular filtration rate (GFR) will continue to decline, eventually necessitating kidney replacement therapy (dialysis or transplant)^8^. The cost of dialysis is high, both in terms of monetary spending ($92-108K per patient each year)^9^ and in terms of quality of life. Thus, there is an urgent need for innovations which elevate GFR and delay or prevent dialysis, improving CKD outcomes.

Changes in gut microbes have been reported in various diseases and conditions, including obesity, diabetes, blood pressure regulation, immune function, and depression^10–13^. Recently, interest has been growing regarding a potential role for the gut microbiota to modulate outcomes in kidney disease^14^. Gut microbial diversity is decreased in CKD patients, and genera *Enterobacteriaceae* is enriched while *Roseburia* is reduced^15–17^. A recent publication reported that CKD patients have significant depletion of *Faecalibacterium*, and that supplementation of *F. prausnitzii* to CKD mice reduces renal dysfunction^18^. Another study reported that gut microbiota depletion attenuates the acute kidney injury (AKI) to CKD transition^19^. We previously noted that amoxicillin treatment after severe ischemic AKI in mice modulates gut microbes to accelerate kidney functional recovery and decrease kidney fibrosis^20^. In this study, we describe a role for gut microbes to modulate GFR in health, as well as in a chronic kidney disease model, suggesting a future possibility of modulating the gut microbiome to support GFR in a clinical setting.

## Methods

### Ethical approval

All animal protocols and procedures were approved by the Johns Hopkins University Institutional Animal Care and Use Committee (accredited by the Association for Assessment and Accreditation of Laboratory Animal Care International).

### Animals

Gut microbiota biomass was depleted using a modified antibiotic treatment protocol with a mixture of 1g/L ampicillin, 1g/L neomycin, and 0.5 g/L vancomycin (ABX, modified from Rakoff-Nahoum et al. 2004) as we have done previously^21^. We previously established that this ABX protocol dramatically suppresses gut microbes within 24 hours, and that this suppression is sustained for weeks^21^. C57BL/6J male and female mice (6-week-old, ordered from Jackson Laboratories, n=7-8) were treated with ABX in drinking water for a total of nine weeks. Glomerular filtration rate (GFR) was measured on weeks 0, 5, and 9 of treatment.

Germ-free (GF) C57BL/6J mice (6-week-old) were randomly assigned into two groups (n=8 per sex per group). One group was continuously housed for 5 weeks as GF mice; the other group was given one oral gavage of a fecal slurry to repopulate microbes (donor mice: n=3 males and females, 4∼5-week-old), and then housed for 5 weeks as conventionalized GF (CGF) mice. We also utilized a control group of age-matched conventional mice (control, born and raised with naturally occurring microbiota, bred at the Johns Hopkins mouse facility). At the end of the week 5, GFR was measured in conventional, GF, and CGF mice.

For experiments modeling CKD, we utilized 0.2% adenine diet to induce CKD in mice, as previously reported^22^. Briefly, C57BL/6J male and female mice (8 weeks old, ordered from Jackson Laboratories) were randomly assigned to four groups (n=8 per sex per group): Chow diet (Research Diets, Cat. No.2018 S), Chow + ABX, Adenine diet (0.2% in chow diet, Research Diets, Cat. No. TD. 220605), Adenine + ABX. The mice were treated with different diets and/or ABX in drinking water for six weeks for females and two weeks for males.

A subset of mice was also given high fat diet. For these experiments, C57BL/6J male and female mice (6-week-old, ordered from Jackson Laboratories) were randomly assigned to four groups (n=8 per sex per group): control diet (CD, Research Diets, Cat. No. D12450J), high fat diet (HFD, Research Diets, Cat. NO. D12492), CD + ABX, or HFD + ABX. The mice were treated with different diets and/or ABX in drinking water for nine weeks.

For empagliflozin (EMPA) treatment experimental design: C57BL/6J male and female mice (6-week-old, ordered from Jackson Laboratories) were randomly assigned to two groups (n=7-9 per sex per group): ABX, or, ABX+EMPA. The mice were treated with ABX or ABX+EMPA in drinking water for five weeks.

### Glomerular filtration rate (GFR)

GFR measurements were performed in conscious and unrestrained mice, as previously reported^23,24^. Briefly, GFR devices (MediBeacon) were affixed to a shaved patch of skin and secured with surgical tape. FITC-sinistrin was then retro-orbitally injected into the mouse at a dose of 7 mg/100 g BW dissolved in sterile saline (0.9% NaCl). For GFR measurements after 4 or 6 weeks of adenine diet, mice were injected with a lower dose (3.5 mg/100 g BW) of FITC-sinistrin. After the FITC-sinistrin injection, the mouse was placed back into the cage with free access to food and water, and clearance was monitored for 2 h post-injection via fluorescent signal decay. Data were analyzed using MPD Studio (MediBeacon). GFR was calculated as volume per minute by determining the rate constant using the t1/2 of the FITC-sinistrin decay curve as In(2)/t1/2, as well as the extracellular fluid volume (ECVS) of each animal as 14,616.8/100 × BW. GFR was ultimately calculated as GFR (ul/min) = rate constant × calculated ECVS. Details regarding these calculations have been reported previously^23^.

### Glucose tolerance test (GTT) and Insulin tolerance test (ITT)

GTT was carried out as previously reported^25^. Briefly, mice were fasted overnight for 16 h and then fasting glucose was measured at time 0 min using a Roche Accu-Chek Nano glucometer, followed by an intraperitoneal (*i.p.*) injection of 1g/kg body weight (BW) glucose (Cat. NO. G8270, Sigma-Aldrich). Blood glucose level was measured with timepoints of 15, 30, 60, 90, 120, and 150 min post-injection.

In separate experiments, ITT were performed as previously described^25^. In brief, mice were fasted for 2 h and then *i.p.* injected with 0.7 Units of insulin (Cat. NO.12585-014, Gibco) /kg BW. Blood glucose levels were measured before injection and at 15, 30, 60, 90, and 120 min post-injection. Data were plotted as mean ± SEM and area under curves for GTT and ITT were calculated.

### qPCR

Fecal samples (two to three pellets) were collected from the mice after six weeks of treatment (four groups: Chow, Chow+ABX, Adenine, Adenine+ABX). Fecal DNA was isolated using the Fast Stool Isolation Kit (catalog no. 51604, QIAGEN). qPCR was performed using bacterial forward primer (bact1369F: 3’-CGGTGAATACGTTCYCGG-5’) and reverse primer (prok1492R: 3’-GGWTACCTTGTTACGACTT-5’).

### Histology

Drop-fixed kidneys stored in 4% paraformaldehyde (PFA) were embedded in paraffin, and 10 um-sections were stained by Picrosirius red staining (section preparation and staining were performed in Johns Hopkins School of Medicine Histology Core Center). Images were taken using a fluorescence microscope (BZ-X700, Keyence). Picrosirius red staining in each section was quantified by ImageJ.

### Other studies

Blood electrolytes, hematocrit, hemoglobin, creatinine, and urea nitrogen (BUN), and non-fasting glucose level were measured via iStat analysis. Plasma creatinine was measured by a Cobas Mira Plus automated analyzer (Roche, Nurley) using Creatinine reagent set (Pointe Scientific, Canton). Kidney weight and body weight were recorded.

### 16S rRNA microbiome analysis

Fecal samples (two to three pellets) were collected from week 6 treated-mice (four groups: Chow, Chow+ABX, Adenine, Adenine+ABX). Fecal DNA was isolated using the Fast Stool Isolation Kit (catalog no. 51604, QIAGEN). We performed 24 PCR cycles for each sample using the following primers: 319F, CTCCTACGGGAGGCAGCAGT and 806R, GGACTACHVGGGTWTCTAAT. 16S rRNA sequencing (was performed by the Johns Hopkins Transcriptomics and Deep Sequencing Core and analyzed by Resphera Biosciences, as previously^26^. Normalization of observations was performed by rarefaction followed by alpha and beta diversity characterization, differential abundance analysis, and unsupervised clustering of microbial profiles. Statistical comparisons of groups of interest employed the Mann-Whitney test supplemented by Welch’s T-test (after log10 transformation for taxonomic levels), and correction of p-values using the False Discovery Rate.

### Statistical analysis

Data are presented as mean ± standard error of the mean (SEM). Data analysis except for 16S rRNA sequencing data, Tukey’s multiple comparison test was performed on measurements in the same group. Comparison between multiple groups was determined by ANOVA with Tukey’s post hoc analysis for multiple groups. Statistical analysis was performed using Prism 9, GraphPad Software, and significance was determined as *p*<0.05.

## Results

### Gut microbes alter GFR in healthy mice

To determine if gut microbiota play a role to set baseline GFR, we measured GFR before and after mice were treated with ABX in drinking water. In females, GFR was significantly increased after ABX treatment for five and nine weeks (**Fig. 1A**). In males, GFR was increased on week 5 of ABX treatment, and the difference was amplified on week 9 (**Fig. 1B**). GFR was not different between females and males before or after treatment, although females trended towards a higher GFR on week 5 of ABX (p=0.052 by two-way ANOVA). We recognize that ABX could have additional effects on host physiology which are unrelated to the suppression of gut microbes. To address this, we next measured GFR in (1) conventional (Ctrl), (2) germ-free (GF), and (3) conventionalized germ-free (CGF) mice. We found that GFR was significantly increased in GF mice as compared to both Ctrl and CGF mice (in females and males, **Fig. 2**). In addition, GFR showed no differences between Ctrl and CGF mice for both females and males (**Fig. 2**). The increased GFR in GF mice (∼24%) was similar in magnitude to the increase seen in ABX-treated mice (∼25%). Of note, GF females have significantly higher GFR than GF males (p=0.0024 by two-way ANOVA). (**Fig. 2**).

**Figure 1.**
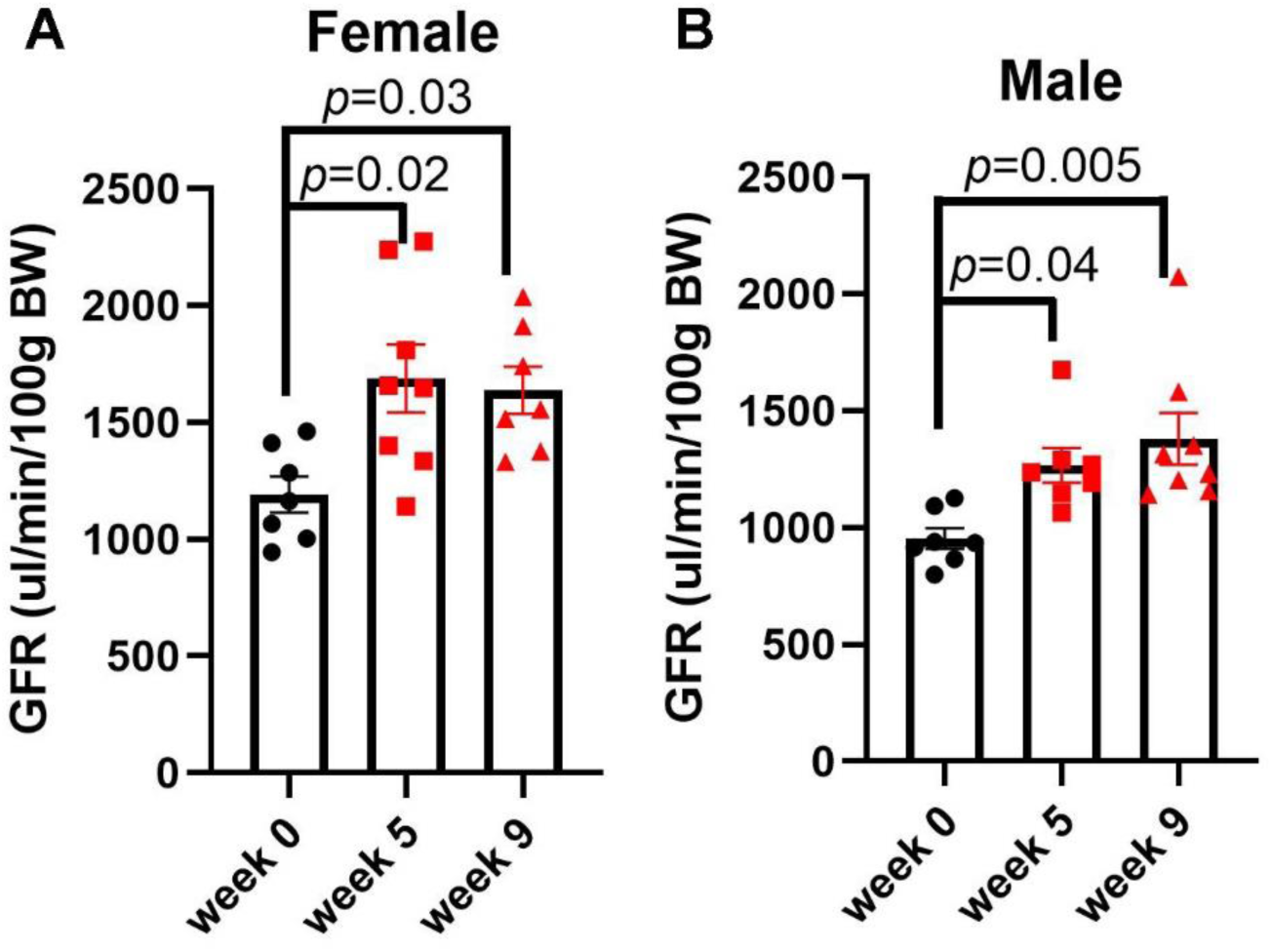
Antibiotic (ABX) treatment increases glomerular filtration rate (GFR) in females and males. GFR was measured before treatment as well as on week 5 and 9 of treatment. GFR was significantly increased after ABX treatment in females **(A)** and males **(B)**. Data presented as mean ± SEM. n=7-8 per group, each dot is an individual mouse. Statistical comparisons were performed by one-way ANOVA.

**Figure 2.**
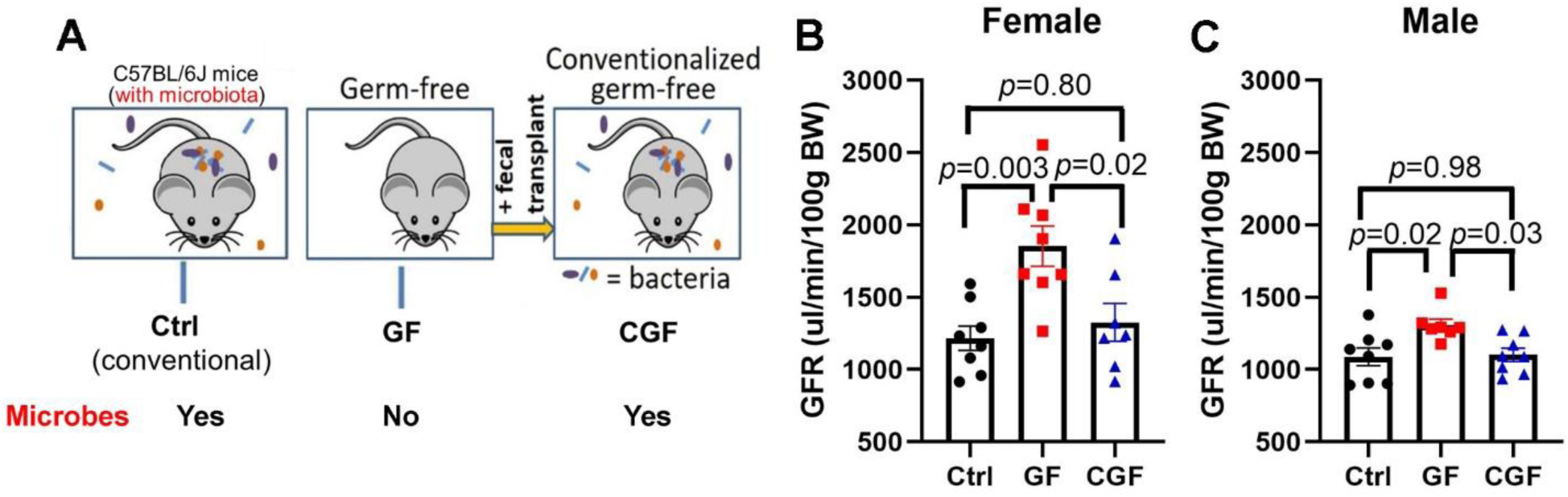
Glomerular filtration rate (GFR) is increased in germ-free (GF) females and males. **(A)** Experimental design. 6-week-old C57BL/6J GF mice were randomly assigned to remain GF, or to be conventionalized with gut microbes (CGF, oral gavaged with fecal slurry). Five weeks later, GFR was measured and compared to age-matched conventional mice. GFR was significantly increased in GF compared to Ctrl and CGF in females **(B)** and males **(C)**. Data presented as mean ± SEM. n=7-8 per group, each dot is an individual mouse. Statistical comparisons were performed by one-way ANOVA.

### Gut microbes alter GFR in a model of CKD in females

The data in Figures 1-2 show that gut microbes alter GFR in healthy mice. To elucidate if gut microbes alter GFR in a chronic kidney disease model, 9-week old mice were randomly assigned into four groups: Chow, Chow+ABX, Adenine, or Adenine+ABX. Mice were treated with different diets and/or with ABX in drinking water for 6 weeks. We hypothesized that adenine treatment would lower GFR, and that this effect would mitigated by ABX. Bacterial suppression was validated by qPCR (**Fig. S1**). In females, GFR was measured on weeks 0, 4 and 6 of treatment (**Fig. 3A-B**). As expected, ABX alone increased GFR and adenine alone decreased GFR on week 4 and week 6. The combination of adenine and ABX improved GFR versus adenine alone on both weeks 4 and 6 of treatment.

**Figure 3.**
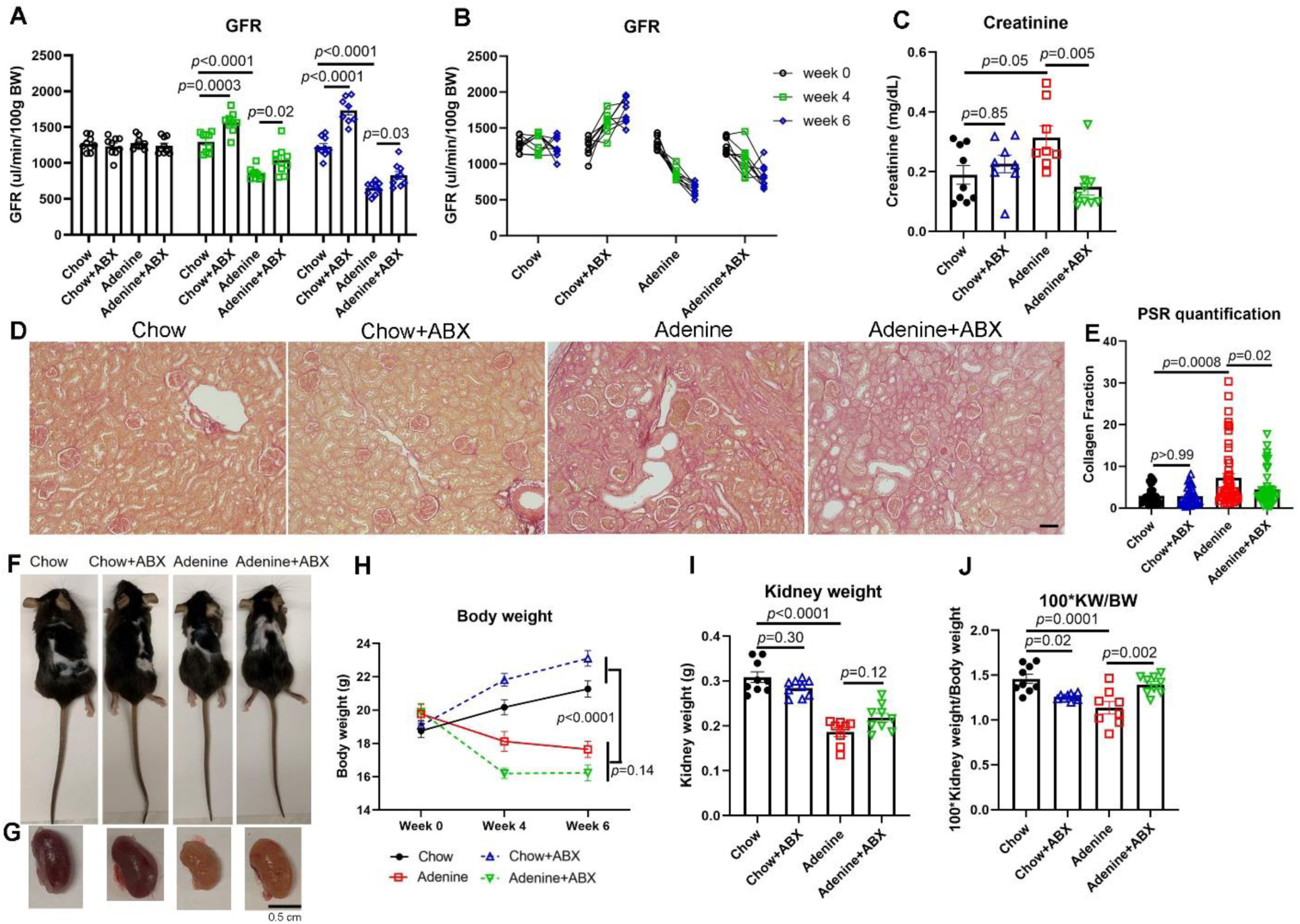
ABX improves kidney function in a model of chronic kidney disease in females. **(A-B)** Adenine treatment alone significantly decreased GFR on week 4 and 6; co-treatment with adenine and ABX delayed the decrease. GFR data from A re-plotted to show trends for each individual mouse; data from the same animal on different weeks were connected with a line. **(C)** Plasma creatinine (on week 6 of treatment) was significantly elevated on adenine diet; this was normalized by ABX treatment. Picrosirius red staining (PSR) was used to stain collagen and evaluate kidney fibrosis on Chow, Chow+ABX, Adenine, Adenine+ABX treatments. ABX did not affect kidney fibrosis on chow diet, however, ABX suppressed the increase in fibrosis seen with adenine diet. Representative images are shown **(D)** and quantification of PSR staining is shown in **(E).** Representative images of mice and kidneys from each group are shown in **(F)** and **(G)**. **(H)** Body weight was significantly decreased on adenine vs chow diet. ABX treatment further lowered body weight when combined with adenine diet. Kidney weight **(I)** was reduced on adenine diet, but ABX did not significantly alter kidney weight on chow or adenine diet. **(J)** The ratio of kidney weight to body weight was increased with adenine + ABX compared to adenine alone. Data are presented as mean ± SEM. Each dot represents one mouse in A-C, I, and J. For D, scale bar: 50 μm. For E, n=30-60 fields of view from n=4∼5 mice per group. n=7-9 per group. Statistical comparisons were performed by one-way or two-way ANOVA.

To provide context to the GFR data, we also measured a variety of other parameters. Plasma creatinine was significantly increased on adenine diet, and was normalized by ABX treatment (**Fig. 3C**). Histological studies revealed that adenine treatment significantly increased collagen, as indicated by picrosirius red staining **(Fig. 3D-E)**. ABX treatment suppressed this increase. Of note, ABX treatment did not affect collagen on chow diet **(Fig. 3D-E)**.

Blood hematocrit and hemoglobin levels were significantly reduced on adenine diet, and ABX treatment did not alter these levels on either chow or adenine diet (**Fig. S2A, B**). BUN was significantly elevated on adenine diet, and ABX treatment did not change BUN on either diet (**Fig. S2C**). Adenine treatment did not alter non-fasting glucose. However, glucose was significantly reduced on adenine+ABX treatment versus chow+ABX treatment (**Fig. S2D**). Adenine treatment decreased the body weight; this was amplified by ABX treatment (**Fig. 3F, H**). However, ABX treatment did not change body weight on chow diet. Kidney weight was reduced by adenine diet, but ABX treatment did not alter kidney weight on either diet (**Fig. 3G, I**). In addition, the ratio of kidney weight to body weight (KW/BW) was significantly decreased on adenine diet (**Fig. 3J**). ABX treatment decreased KW/BW on chow diet, but increased it on adenine diet (**Fig. 3J**).

### Gut microbes alter GFR in a model of CKD in males

A pilot study indicated that the GFR decease with adenine diet began at an earlier timepoint in male mice. In addition, the body weight was decreased more than 20% on week 4 of adenine treatment. Thus, the measurements were performed on week 2 of treatment for males (**Fig. 4A-B**). As expected, ABX alone increased GFR, and adenine alone decreased GFR, the combination of adenine+ABX improved GFR.

**Figure 4.**
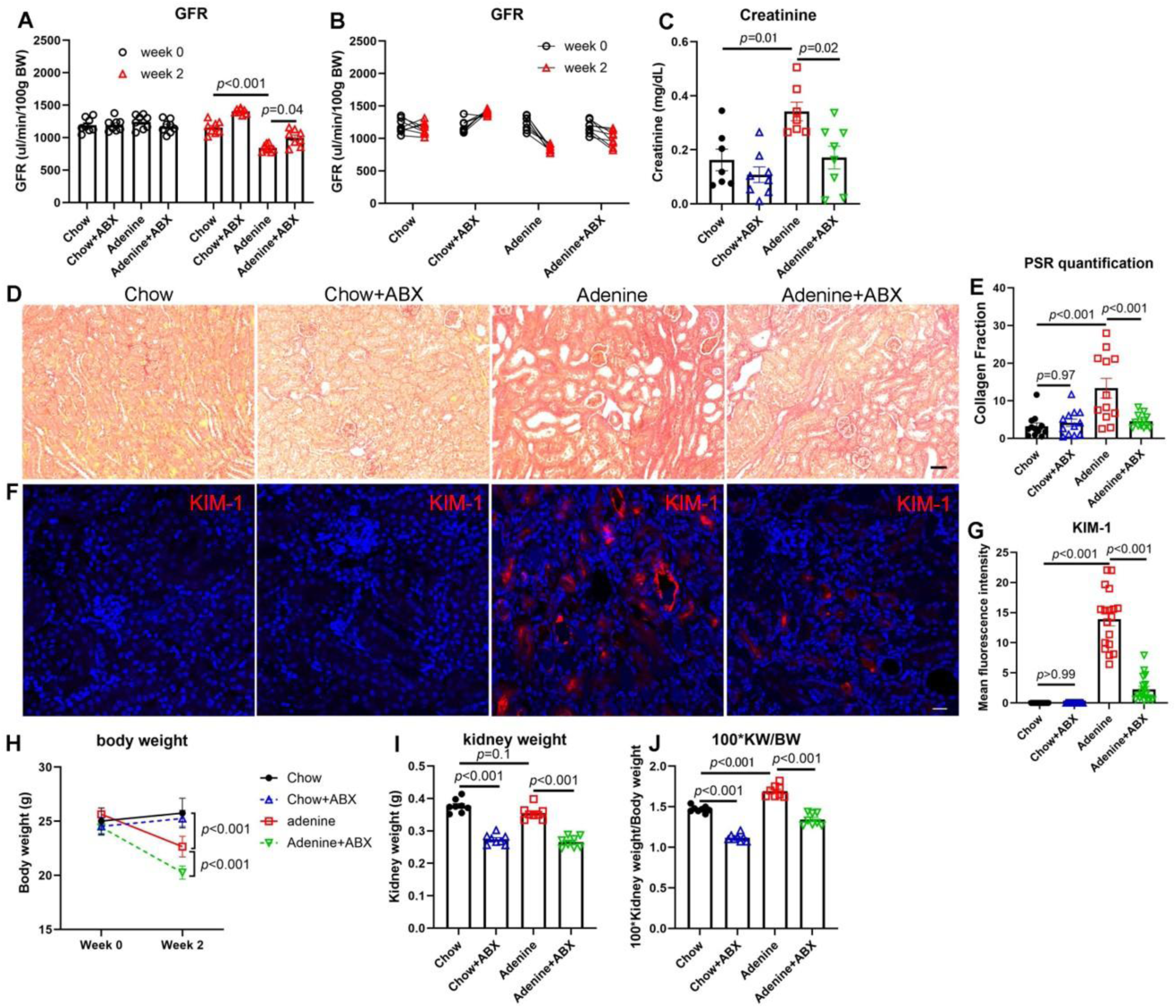
ABX improves kidney function in a model of chronic kidney disease in males. **(A-B)** Adenine treatment alone significantly decreased GFR on week 2; co-treatment with adenine and ABX delayed the decrease in GFR. GFR data from A re-plotted to show trends for each individual mouse; data from the same animal on different weeks were connected with a line. **(C)** Plasma creatinine was significantly elevated on adenine diet; this was partially mitigated by ABX treatment. Picrosirius red staining (PSR) was used to stain collagen and evaluate kidney fibrosis on Chow, Chow+ABX, Adenine, and Adenine+ABX treatments. Representative images are shown in **(D)**, and quantification is shown in **(E).** ABX did not affect kidney fibrosis on chow diet. However, ABX suppressed the increase in fibrosis seen with adenine diet. Adenine treatment significantly increased kidney injury molecule (KIM-1) which was reduced by ABX. Representative images are shown in **(F)** and quantification is in **(G)**. **(H)** Body weight was significantly decreased on adenine vs chow diet. **(I)** ABX treatment further lowered body weight when combined with adenine diet. Adenine diet did not alter kidney weight, but ABX significantly decreased kidney weight on either diet. **(J)** The ratio of kidney weight to body weight was increased with adenine, but ABX reduced the ratio on either diet. Data are presented as mean ± SEM. Each dot represents one mouse in A-C, I-J, n=7-9 per group. For D, scale bar: 50 μm. n=12 kidney sections from n=4 mice per group. For F, scale bar: 20 μm. n=15∼25 fields of view from n=3 mice per group. Statistical comparisons were performed by one-way or three-way ANOVA.

Plasma creatinine was significantly elevated with adenine diet, but co-treatment with ABX was protective (**Fig. 4C**). Picrosirius red staining indicated that collagen was significantly increased on adenine diet which was suppressed by ABX treatment **(Fig. 4D-E)**. Of note, ABX treatment did not affect collagen on chow diet **(Fig. 4D-E)**. Further analysis focused on kidney injury molecule (KIM-1) indicated that adenine treatment significantly increased KIM-1 which was reduced by ABX **(Fig. 4F-G)**.

Body weight was decreased on adenine diet; this decrease was exacerbated by ABX treatment (**Fig. 4H**). ABX treatment did not alter body weight on chow diet. Adenine diet did not alter kidney weight, but ABX significantly decreased kidney weight on either diet (**Fig. 4I**). The ratio of kidney weight to body weight (KW/BW) was increased with adenine vs chow diet, but ABX reduced the ratio on either diet (**Fig. 4J**).

Adenine treatment slightly reduced blood hematocrit and hemoglobin levels, but ABX treatment did not alter these levels on either diet (**Fig. S2E, F**). Blood analysis demonstrated that BUN was significantly increased on adenine than chow diet mice, but was not altered by ABX on either diet (**Fig. S2G**). Adenine treatment also decreased non-fasting glucose (**Fig. S2H**) and the combination of adenine with ABX further reduced the glucose level.

### A potential role for *Akkermansia muciliphila* in GFR regulation

These data imply that gut microbiota produce signals which can alter GFR in both health and disease. To identify bacterial species which may contribute to GFR regulation in adenine-induced CKD, we analyzed fecal 16S ribosomal RNA (rRNA) from female mice treated for 6 weeks with Chow, Chow+ABX, Adenine, or Adenine+ABX (n=6 per group). The relative abundances of phyla among these four groups is shown in **Fig. 5A**. Adenine treatment significantly increased *Verrucomicrobia* compared to chow diet, which was dramatically reduced by ABX (**Fig. 5A-B**). Further analysis indicated that *Akkermansia muciliphila* drove the elevation of *Verrucomicrobia* phylum with adenine, and was suppressed by the combination of adenine and ABX (**Fig. 5C**, Adenine vs Adenine+ABX, *p*=0.04 via Mann-Whitney). These data suggest that *Akkermansia muciliphila* plays a potential role in GFR regulation in adenine-induced CKD females.

**Figure 5.**
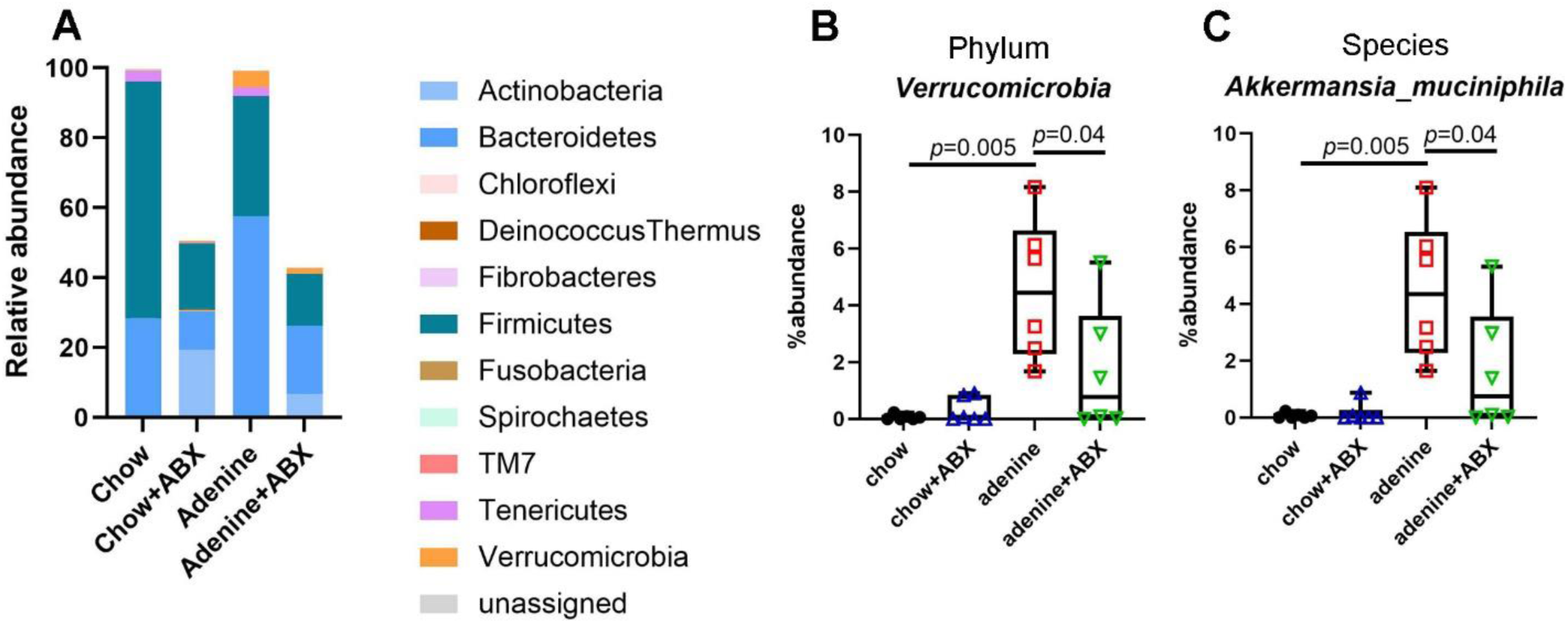
ABX treatment alter gut microbial abundance in adenine-induced CKD females. **(A)** Relative abundance of gut microbiome at the phylum level. **(B)** Compared to chow diet, adenine treatment was significantly increased *Verrucomicrobia* (*p*=0.005) which was reduced by ABX (*p*=0.04). **(C)** *Akkermansia muciniphila* under *Verrucomicrobia* phylum was significantly increased (*p*=0.005, adjusted *p* value *p*=0.21) in adenine and decreased by the combination of adenine and ABX (*p*=0.04; adjusted *p* value *p*=0.17). Each column represents a sample from a different animal (F1, F2, etc). n=6 per group. Statistical comparisons were performed by the Mann-Whitney test (\**p*<0.05, \*\**p*<0.01).

### Increased GFR seen with HFD and ABX is additive in females but not in males

In healthy mice, we observed that ABX induce an elevation of GFR above baseline (i.e., hyperfiltration). Hyperfiltration can also occur in diabetes, and we^23^ and others^27,28^ have shown that high fat diet (HFD) is sufficient to induce hyperfiltration in C57BL6/J mice. Thus, to determine if there may be a common mechanism, we asked whether the increase in GFR seen with ABX and HFD is additive. To examine this, 6-week-old C57BL/6J females and males were randomly assigned into four groups: control diet (CD), CD+ABX, HFD, or HFD+ABX. Mice were treated with different diets and/or with ABX in drinking water for 10 weeks. HFD alone and ABX alone both increased GFR at week 5 and week 9 in both sexes (**Fig. 6**). Co-treatment with HFD and ABX resulted in a further increase in GFR in females at both week 5 and week 9 (**Fig. 6A, C**); however, there was no additional GFR increase with co-treatment in males (**Fig. 6B, D**). Of note, HFD males had significantly higher GFR than HFD females on week 9 of treatment (*p*=0.015 via two-way ANOVA).

**Figure 6.**
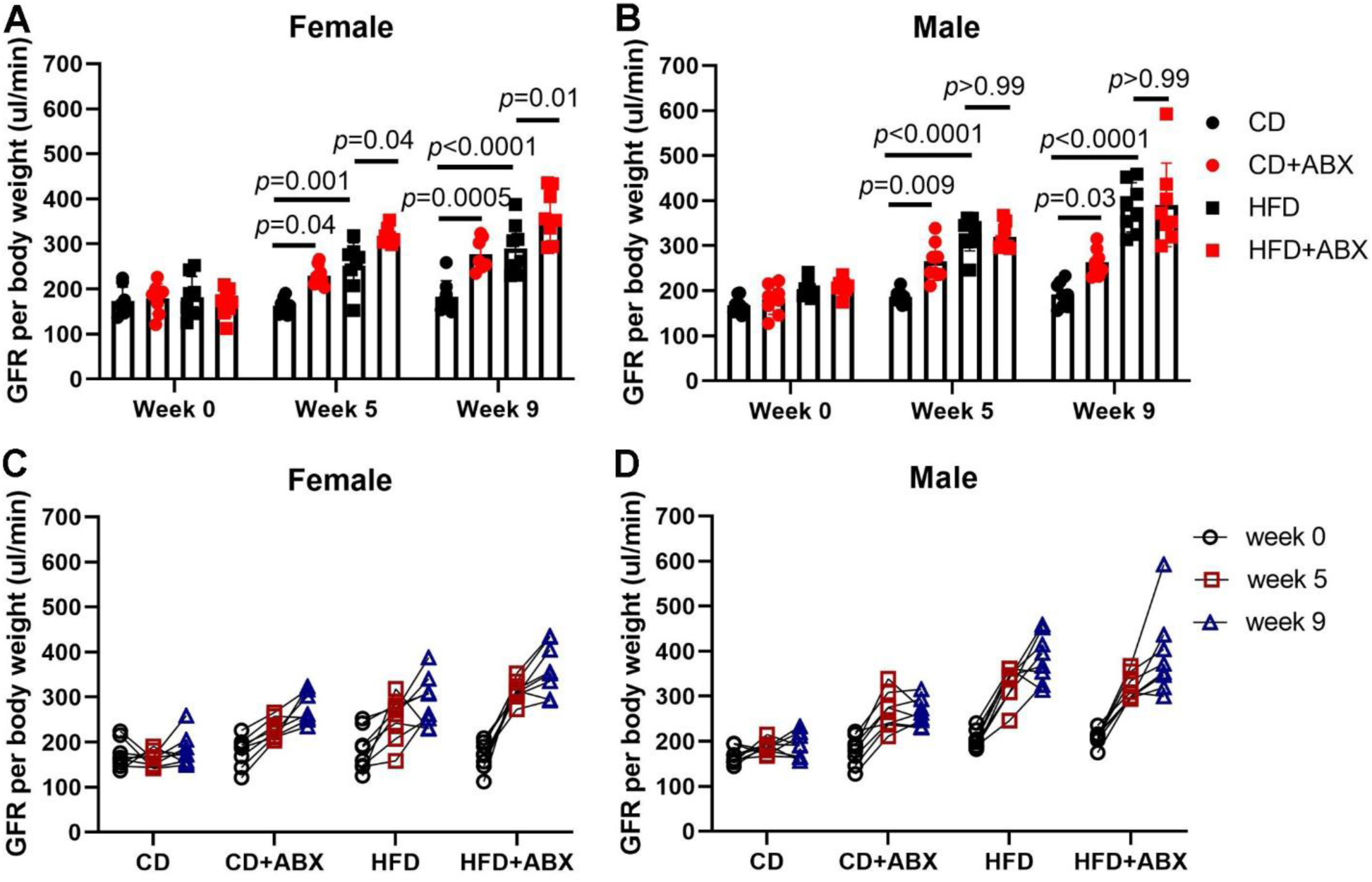
Both high fat diet (HFD) and antibiotics (ABX) increase GFR; these effects are synergistic in females. **(A)** ABX significantly increased GFR on both control diet (CD) and HFD in females on weeks 5 and 9. **(B)** ABX treatment elevated GFR only on CD but not on HFD in males. GFR data from A-B are replotted in **(C)** and **(D)** to show trends for individual animals. The same mouse on different weeks was connected with a line. Data are presented as mean ± SEM. Each dot represents one mouse in **A** and **B**. n=6-8 per group. Statistical comparisons were performed by three-way ANOVA.

We observed no differences in blood electrolytes, glucose, hematocrit, hemoglobin, creatinine, or BUN in mice treated with vs without antibiotics on either data (in both females and males; **Table S1 and S2**); of note, creatinine here was measured by the less-sensitive iStat (as compared to data in Figs 3-4). Although HFD altered body weight, GTT responses, and plasma insulin levels, ABX treatment did not (**Fig. S3-S5**). However, we did observe a decrease in the area under the curve (AUC) for ITT in mice on control diet treated with ABX.

### Evaluating a role for tubuloglomerular feedback (TGF) in microbial control of GFR

The mechanism of GFR increase with HFD is thought to involve alternations in tubuloglomerular feedback (TGF)^29^. We hypothesized that something similar may happen with ABX treatment: if Na^+^ reabsorption in proximal tubule were to be elevated when microbial signals are absent or suppressed, the reduced Na^+^ delivery to the macula densa would alter TGF to increase GFR. To test this, we used the sodium-glucose cotransporter 2 (SGLT2) inhibitor empagliflozin (EMPA) which impairs Na^+^ reabsorption in proximal tubule, increasing Na^+^ delivery to the macula densa. Mice were randomly assigned to two groups: ABX or ABX+EMPA (ABX with or without EMPA was given in drinking water for 5 weeks; **Fig. 7A**). As expected, GFR was significantly increased after ABX treatment at 3 and 5 weeks in both females and males. The combination of ABX and EMPA treatment resulted in a delayed elevation of GFR (elevated GFR was seen at week 5 but not week 3, **Fig. 7B, C**). These data suggest that EMPA partially inhibits microbial regulation of GFR in both females and males. There was no sex difference in GFR at week 0, but females had significantly higher GFR on week 5 on ABX (*p*=0.016 via two-way ANOVA). Sex differences were not observed between males and females treated with both EMPA and ABX.

**Figure 7.**
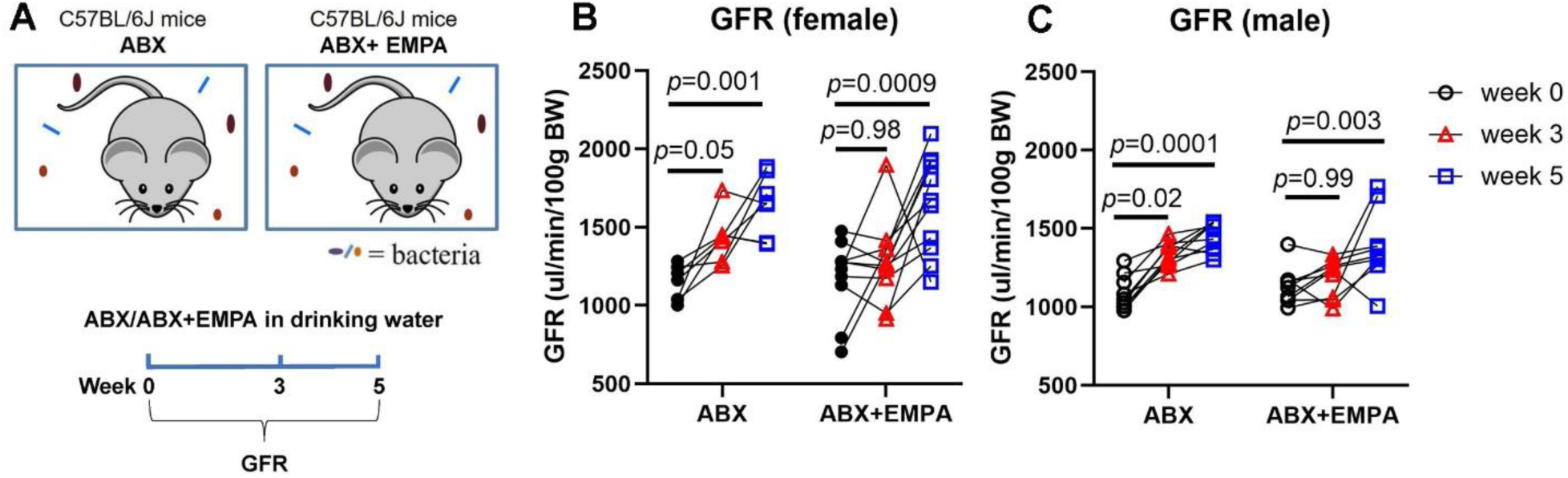
EMPA partially reduces ABX-induced GFR increases on both sexes. **(A)** Experimental design. Females and males were treated with ABX or ABX + EMPA for 5 weeks in total. EMPA treatment normalized ABX-induced GFR increases on week 3 but not on week 5 in females **(B)** and males **(C)**. n=7-9 per group. Statistical comparisons were performed by ANOVA.

## Discussion

In this study, we identified a role for the microbiome to modulate GFR in murine health and chronic kidney disease. We report that ABX treatment increased GFR by ∼25% in healthy mice; similarly, we find that GF mice had an ∼25% elevation in GFR. In a CKD model with impaired GFR, we likewise found that the adenine-induced decrease in GFR was attenuated by ABX. Mice treated with both adenine and ABX also had improvements in plasma creatinine and renal fibrosis as compared to mice treated with adenine alone, and we report that adenine diet induces dramatic changes in *Akkermansia muciliphila*. However, animals treated with both adenine and ABX had a worsened weight loss. To explore potential mechanisms by which microbes modulate GFR, we utilized mice treated with HFD as well as mice treated with an SGLT2 inhibitor. These studies showed that altered TGF likely contributes to – but does not fully explain – the microbial modulation of GFR.

### Microbiome and GFR in health

To our knowledge, the relationship between the microbiome and GFR in health has not been investigated previously. We find that GFR is increased in ABX-treated (microbiome suppressed) and GF (microbiome absent) females and males, suggesting that microbes play an essential role to ‘set’ baseline GFR. These data imply that GFR is chronically held in check by microbial signals, and that in the absence of such signals, the inhibition is released and GFR is elevated. Although the identity of such signals is unknown, it is tempting to hypothesize that gut microbial metabolites may play a role.

### Microbiome and GFR in CKD

To elucidate if microbes regulate GFR in a CKD model where GFR is impaired, we measured GFR in adenine-induced CKD mice. ABX improved GFR in adenine treated females on both weeks 4 and 6, and, in adenine treated males on week 2. These data suggest that the microbiome modulates GFR in this CKD model. Similarly, plasma creatinine was increased with adenine treatment but reduced by co-treatment with ABX. Of note, the lower body weight in Adenine+ABX mice may reflect a lower muscle mass, which could also affect plasma creatinine. Our findings are consistent with previous reports that the gut microbiota is correlated with kidney function in disease. For example, it was reported that microbiome diversity is associated with estimated GFR in CKD patients^15–17,30–33^. Another study reported that microbiota from end-stage renal disease patients transplanted to renal injured GF mice or antibiotic-treated rats aggravated renal fibrosis and oxidative stress^34^. The same study further indicated that two species, *Eggerthella lenta* and *Fusobacterium nucleatum,* increase uremic toxin production and promote CKD development, whereas the probiotic *Bifidobacterium animalis* reduces abundance of these species, leading to lower toxin levels and improving disease^34^. A separate study reported that the prebiotic *gum acacia* had a beneficial role in CKD^35^. Of note, the microbiome is not only associated with GFR in CKD, but also with acute kidney injury (AKI): we previously reported that modification of gut microbiota after severe ischemic AKI accelerates kidney recovery and mitigates the progression of AKI to CKD^24^.

In our study, we used 16S rRNA analysis to show that *Akkermansia muciliphila* was increased with adenine diet. Our finding aligns with a report that 5/6 nephrectomy C57BL/6J mice have a higher relative abundance of *Akkermansia muciniphila*^36^. In addition, another study also reported that *Akkermansia* is enriched in adenine-induced CKD rats^37^. However, studies on CKD patients are conflicting: some reported that the abundance of Akkermansia increased along with the progression of CKD^16,38,39^, but others found that *Akkermansia muciniphila* is reduced in CKD patients^40^.

### Potential mechanisms by which the microbiome modulates GFR

Elevations in GFR above baseline are fairly uncommon, but it is known that GFR is significantly elevated in high fat diet-induced hyperfiltration^27^ through a mechanism involving TGF^41^. A previous report used EMPA to disrupt TGF and reduced renal hyperfiltration in a unilateral nephrectomy mouse model^42^. We treated ABX-treated mice with the EMPA to disrupt TGF, and found that EMPA treatment restores GFR to normal levels in ABX-treated mice at early timepoints, but not at later timepoints. Although we did not measure GFR in control (no ABX) mice with and without EMPA, previous studies have reported that GFR was not altered in C57BL/6J mice either treated with 150 mg/kg EMPA of diet for 16 weeks^43^ or 30 mg/kg/day EMPA for 2 weeks^42^. Single nephron GFR studies likewise indicated that EMPA had no effect on GFR in healthy C57BL/6 mice treated with 20 mg/kg/day EMPA by oral gavage for 4 weeks^44^. In addition, EMPA (10 mg/kg/day for 4 weeks) did not change GFR in healthy microbiome intact rats^45^. Although we used EMPA in our study as a pharmacological inhibitor of SGLT2, we should acknowledge that a recent study reported that dapagliflozin (a different SGLT2 inhibitor) alters the abundance of bacteria taxa and decreases uremic toxins^46^. Similarly, the SGLT2 inhibitor canagliflozin contributes to reconstructing the intestinal flora and decreases uremic toxins in Dahl salt-sensitive rats^47^. Thus, we cannot rule out a more direct microbial effect of EMPA in our studies, in addition to inhibiting SGLT2 itself, which could complicate interpretation of our data. Nevertheless, these data imply that EMPA treatment can delay the effect of ABX on GFR.

It should be noted that immune responses may also play a role in the microbiome effect on GFR. Although not investigated in this study, we previously reported that microbiome modulation improved functional recovery after acute kidney injury, and that this effect was dependent on CD8^+^T lymphocytes^20^. Thus, in the future it will be important to determine whether the effects seen in this study may be partially mediated by changes in immune cells.

Another key area for future study is a potential role for microbial metabolites in mediating the gut microbial effect on GFR. Evidence suggests that, although GF mice have significantly lower levels of uremic toxins, they experience more severe renal damage compared to specific pathogen-free mice with adenine-induced renal failure, likely due to the loss of renoprotective short-chain fatty acids^48^. Previous studies have found that microbial effects on host phenotypes can be driven by changes in circulating metabolites, which then interact with host receptors or pathways^18,49–51^. Studies focused on metabolites related to GFR in CKD patients found that some metabolites are positively but some are negatively associated with estimated GFR^52–55^. Of note, the secretion of microbial metabolites can be influenced by transporters such as organic anion transporter-1 (OAT-1)^56^. Thus, one possibility is that the influence of the gut microbiome on GFR is driven by changes in circulating levels of gut microbial metabolites, which will be a major focus of the future mechanistic studies

In sum, we find that the gut microbiome plays a role to establish the GFR setpoint in healthy mice, and that suppressing gut microbes can elevate GFR in both health and chronic kidney disease. In future studies, it will be important to further explore how TGF and other mechanisms may contribute to this regulation, and to further examine the role of specific microbes such *Akkermansia muciniphila* in GFR modulation.

## List of abbreviations

GFR: glomerular filtration rate
CKD: chronic kidney disease
ABX: antibiotics
GF: germ free
CGF: conventionalized germ free
CD: control diet
HFD: high fat diet
EMPA: empagliflozin
GTT: glucose tolerance test
ITT: insulin tolerance test
PFA: paraformaldehyde
BUN: blood urea nitrogen
AUC: area under curve
TGF: tubuloglomerular feedback
SGLT2: sodium-glucose cotransporter 2

## Acknowledgements

We thank past and present Pluznick laboratory members for helpful discussion, suggestions, and comments. We thank Jasmeet Sethi for performing 16S rRNA sequencing.

## Authors’ Contributions

J.L.P., J.X. and H. R. conceived and designed the research. J.X., E.V., J.S., K.G., S.G., and M.G. performed the experiments. J.L.P., J.X. and H. R. interpreted the data. J.X. and J.L.P. wrote the manuscript. J.L.P. and J.X. provided funding. All authors read and approved the final manuscript.

## Data Availability Statement

All data needed to evaluate the conclusions in the paper are present in the paper and the supplementary materials. 16S rRNA sequencing data were uploaded to NCBI SRA (the link is pending and will be provided during the revision).

