## supplementary figures for "Microbes regulate glomerular filtration rate in health and chronic kidney disease in mice"

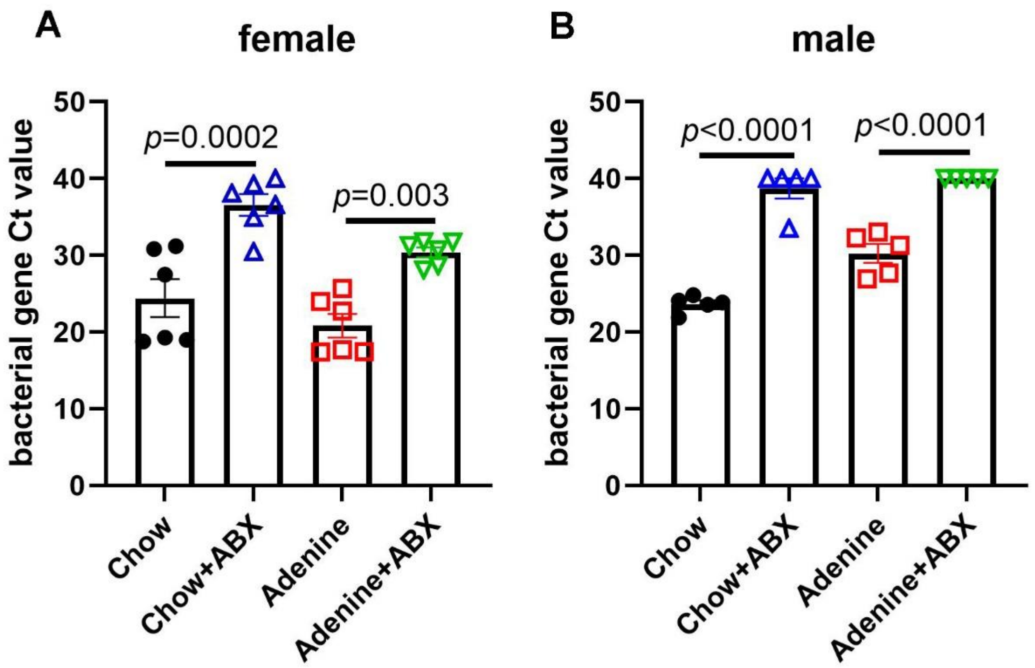


**Figure S1. Bacterial gene reduction after antibiotics (ABX) treatment on both Chow and Adenine diet.** Females and males were treated with chow or adenine diet and/or ABX for 6 weeks, feces DNA were extracted for qPCR. The Ct values for bacterial gene was dramatically reduced after ABX treatment in both females **(A)** and males **(B)**. Each dot represents one mouse. n=5-6 per group. Statistical analysis was performed one-way ANOVA.

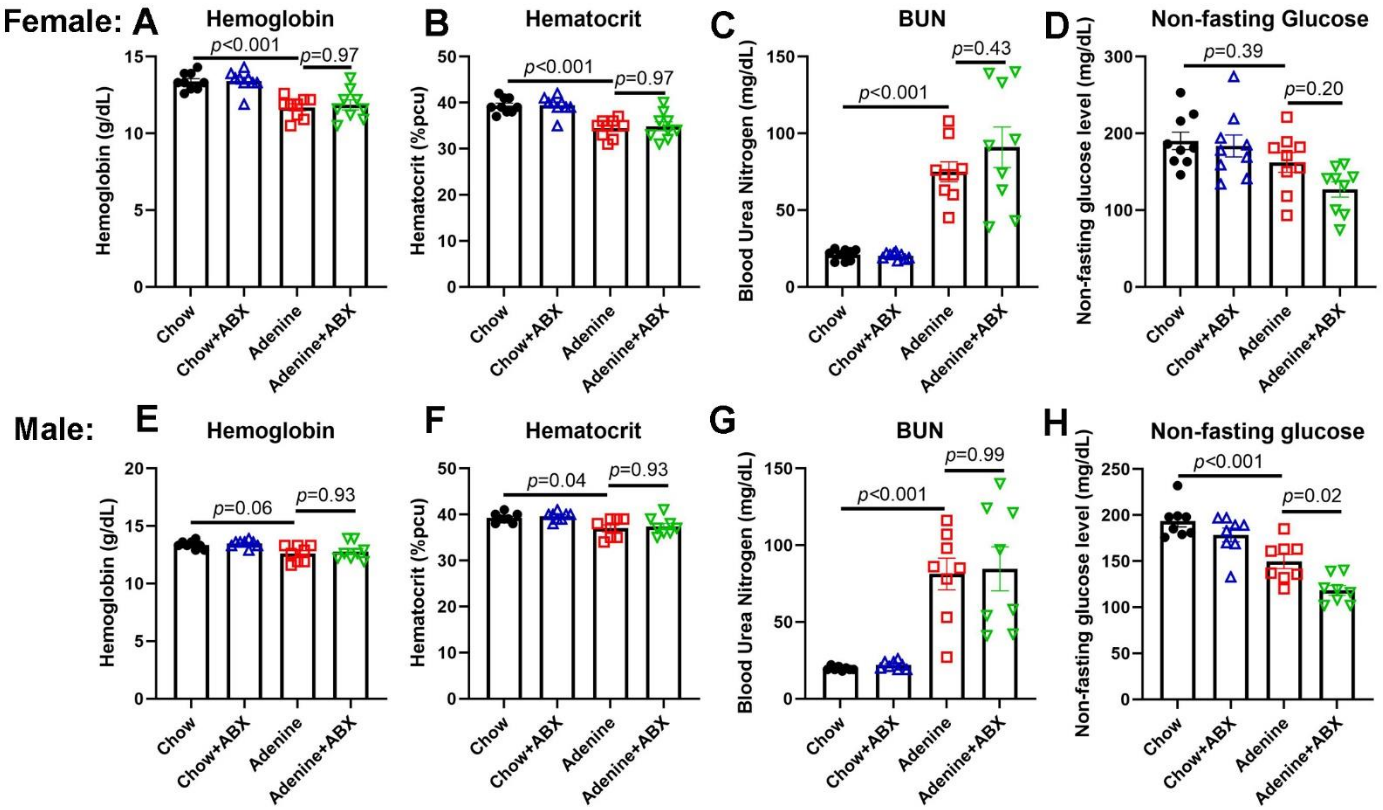


**Figure S2. Blood analysis on CKD females and males.** Blood was analyzed via iStat system after 6 weeks of adenine and/or ABX treatment for females, and after 2 weeks for males. BUN: blood urea nitrogen. Each dot represents one mouse. n=7-9 per group. Statistical analysis was performed one-way ANOVA.

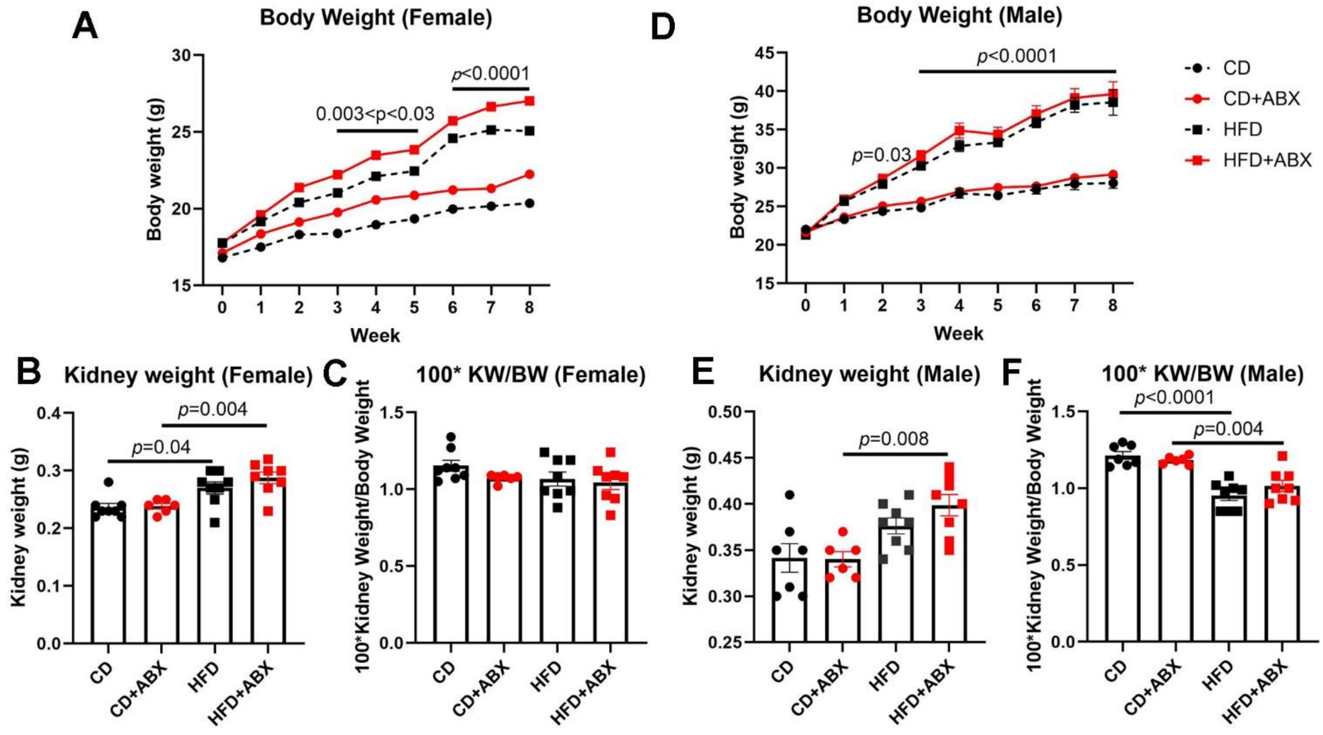


**Figure S3. ABX treatment does not alter body weight or kidney weight on control diet (CD) or high fat diet (HFD).** In females, high fat diet (HFD) increased body weight (BW) **(A)** and kidney weight (KW) **(B)** but did not affect KW/BW **(C)**. In males, HFD increased BW **(D)** and the combination of HFD and ABX treatment increased KW versus CD+ABX **(E)**. HFD reduced KW/BW in males **(F)**. CD: chow diet; HFD: high fat diet; ABX: antibiotics. Data are mean ± SEM. Each dot represents one mouse in **B-F**. n=6-8 per group. Statistical analysis was performed one-way ANOVA.

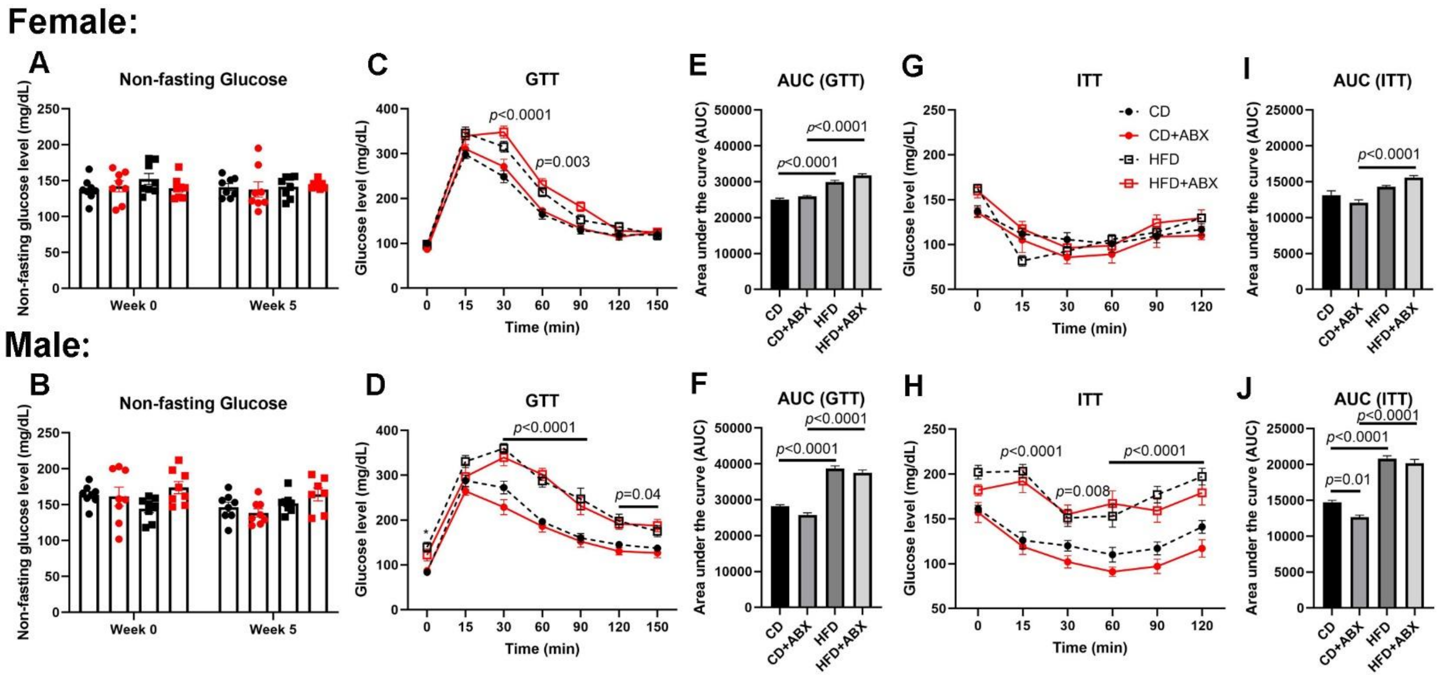


**Figure S4. ABX treatment does not alter GTT or ITT.** HFD or ABX treatment did not alter non-fasting glucose in either females or males **(A-B)**. Glucose tolerance tests (GTT) and insulin tolerance tests (ITT) were altered by HFD, but, not by ABX treatment **(C-J)**. CD: chow diet; HFD: high fat diet; ABX: antibiotics. AUC = area under the curve. Data are mean ± SEM. Each dot represents one mouse in **A-B**. n=6-8 per group. Statistical analysis was performed by one-way ANOVA.

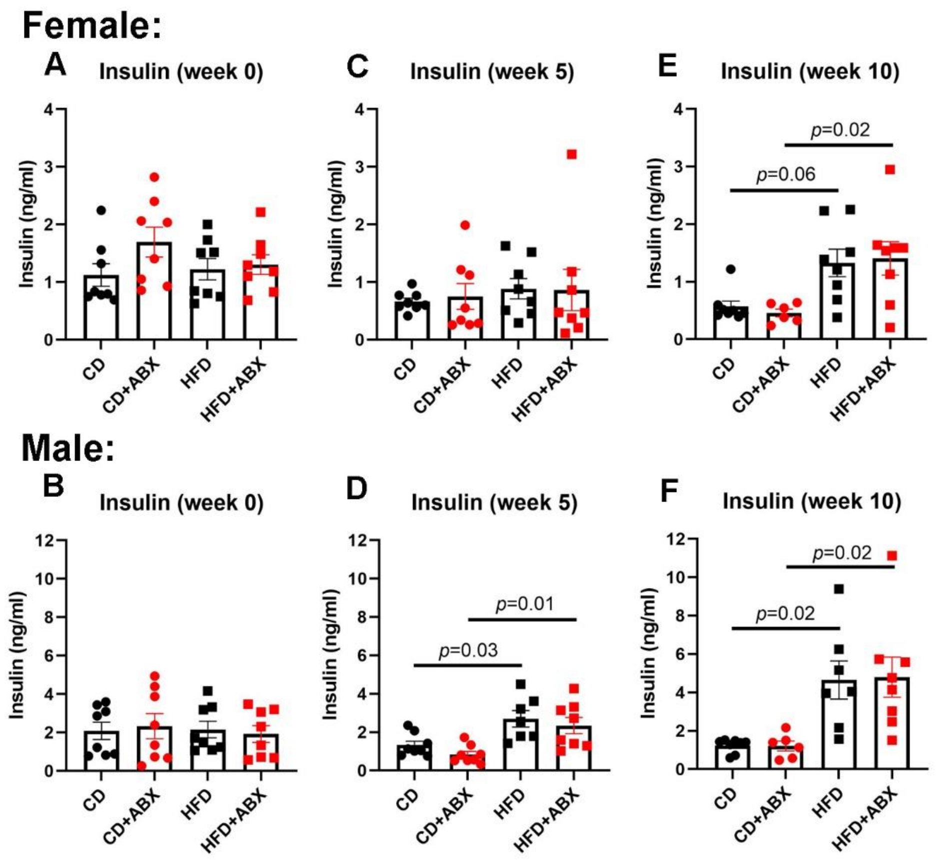


**Figure S5. ABX treatment does not alter Plasma insulin level.** ABX treatment did not alter plasma insulin in females or males **(A-F)**. HFD increased insulin at week 5 for males **(E)**, and at week 10 for both males and females **(C, F)**. CD: chow diet; HFD: high fat diet; ABX: antibiotics. Data are mean ± SEM. Each dot represents one mouse. n=6-8 per group. Statistical analysis was performed by one-way ANOVA.

**Table S1. Blood electrolytes in control diet (CD) or high fat diet (HFD) with or without antibiotics (ABX)-treated females**

| | CD, F | CD+ABX, F | HFD, F | HFD+ ABX, F | *p* (CD vs ABX) | *p* (HFD vs ABX) |
|---|---|---|---|---|---|---|
| **Na, mM/L** | 146.3±0.48 | 147.0±0.1 | 146.3±0.25 | 145.0±0.71 | 0.35 | 0.15 |
| **Cl, mM/L** | 110.3±0.25 | 109.0±0.2 | 113.0±0.41 | 110.8±0.48 | 0.16 | 0.10 |
| **iCa, mM/L** | 1.2±0.05 | 1.2±0.01 | 1.3±0.01 | 1.3±0.01 | 0.89 | 0.11 |
| **BUN, mg/dl** | 21.0±1.08 | 23.5±0.5 | 25.0±1.73 | 21.8±0.95 | 0.83 | 0.77 |
| **Creatinine, mg/dl** | <0.2 | <0.2 | <0.2 | <0.2 | | |
| **Hematocrit, %PCV** | 38.3±3.57 | 42.5±0.5 | 41.3±0.85 | 41.0±1.22 | 0.47 | 0.87 |
| **Hemoglobin, g/dl** | 13.0±1.23 | 14.5±0.15 | 14.0±0.28 | 13.9±0.43 | 0.48 | 0.85 |

Blood chemistries of ABX-treated females on different diets are similar. F: females. iCa: ionized calcium; BUN: blood urea nitrogen; PCV: packed cell volume. The last two columns show p values for a one-way ANOVA for CD vs CD+ABX, or, HFD vs HFD+ABX.

**Table S2. Blood electrolytes in control diet (CD) or high fat diet (HFD) with or without antibiotics (ABX)-treated males**

| | CD, M | CD+ ABX, M | HFD, M | HFD+ ABX, M | *p* (Ctrl vs ABX) | *p* (HF vs ABX) |
|---|---|---|---|---|---|---|
| **Na, mM/L** | 147.8±0.25 | 147.3±0.55 | 146.0±0.58 | 147.3±0.25 | 0.55 | 0.1 |
| **Cl, mM/L** | 112.0±0.41 | 114.5±0.5 | 114.0±0.58 | 114.5±1.19 | 0.1 | 0.72 |
| **iCa, mM/L** | 1.2±0.01 | 1.2±0.03 | 1.2±0.04 | 1.2±0.01 | 0.95 | 0.75 |
| **BUN, mg/dl** | 23.5±0.96 | 20.3±0.85 | 23.3±1.25 | 25.0±1.08 | 0.1 | 0.33 |
| **Creatinine, mg/dl** | <0.2 | <0.2 | <0.2 | <0.2 | | |
| **Hematocrit, %PCV** | 42.8±1.18 | 42.5±1.55 | 39.8±1.31 | 44.3±1.11 | 0.9 | 0.1 |
| **Hemoglobin, g/dl** | 14.5±0.40 | 14.5±0.52 | 13.5±0.46 | 15.1±0.38 | 0.94 | 0.1 |

Blood chemistries of ABX-treated males on different diets are similar. M: males. iCa: ionized calcium; BUN: blood urea nitrogen; PCV: packed cell volume. The last two columns show p values for a one-way ANOVA for CD vs CD+ABX, or, HFD vs HFD+ABX.
